# Screening and Identification of Lassa Virus Endonuclease-targeting Inhibitors from a Fragment-based Drug Development Library

**DOI:** 10.1101/2021.07.09.451867

**Authors:** Xiaohao Lan, Yueli Zhang, Yang Liu, Jiao Guo, Xiaoying Jia, Mengmeng Zhang, Junyuan Cao, Gengfu Xiao, Yu Guo, Wei Wang

**Author notes:** Address correspondence to Wei Wang.

## Abstract

Lassa virus (LASV) belongs to the Old World genus *Mammarenavirus*, family *Arenaviridae*, and order *Bunyavirales*. Arenavirus contains a segmented negative-sense RNA genome, which is in line with the bunyavirus and orthomyxoviruses. The segmented negative-sense RNA viruses utilize a cap-snatching strategy to provide primers cleavaged from the host capped mRNA for viral mRNA transcription. As a similar strategy and the conformational conservation shared with these viruses, the endonuclease (EN) would serve as an attractive target for developing broad-spectrum inhibitors. Using the LASV minigenome (MG) system, we screened a fragment-based drug development library and found three candidates (F1204, F1781, and F1597) inhibited MG activity. All three candidates also inhibited the prototype arenavirus Lymphocytic choriomeningitis virus (LCMV) MG activity. Furthermore, the investigation revealed that two benzotriazole compounds (F1204 and F1781) effectively inhibited authentic LCMV and severe fever with thrombocytopenia syndrome virus (SFTSV) infections. The combination of either compound with an arenavirus entry inhibitor had significant synergistic antiviral effects. Moreover, both F1204 and F1781 were found to exert the binding ability of LASV EN with binding affinity at the micromolar level. These findings provide a basis for developing benzotriazole compounds as potential candidates for the treatment of segmented negative-sense RNA virus infections.

**Importance:** Cap-snatching is the mRNA transcription strategy shared by all the segmented, negative-sense RNA viruses. Using a fragment-based drug development (FBDD) library, we tried to screen out the backbone compound to inhibit the endonuclease activity and thus block this kind of virus infection. Two benzotriazole compounds, F1204 and F1781, were identified to inhibit the Lassa virus (LASV) minigenome activity by targeting the LASV EN.

## Introduction

Lassa virus (LASV) is an enveloped, negative-sense, bi-segmented RNA virus that belongs to the genus *Mammarenavirus* (family *Arenaviridae*) (1). LASV causes the hemorrhagic disease Lassa fever (LF), an annual epidemic in West Africa and peaks during the dry season. In 2020, Nigeria faced a large outbreak of LF, with 1,189 confirmed cases and 244 deaths, according to the Nigeria Centre for Disease Control. As LASV is associated with a high mortality rate in humans, it is listed as a biosafety level 4 (BSL-4) agent. Most pathogenic mammarenaviruses, including the Junín virus (JUNV), Machupo virus (MACV), Guanarito virus (GTOV), Chapare virus (CHAPV), Sabiá virus (SBAV), and Lujo viruses (LUJV), are known to cause severe hemorrhagic fever and are also listed as BSL-4 agents. However, the prototypes of *Mammarenavirus*, Lymphocytic choriomeningitis virus (LCMV), a close relative of LASV, is usually asymptomatic and is categorized as a BSL-2 agent (2).

*Mammarenavirus* RNA genomes contain the S segment, encoding glycoprotein complex (GPC) and nucleoprotein (NP), and the L segment, encoding matrix protein (Z) and viral polymerase (L). In line with other segmented, negative-sense RNA viruses, such as influenza A virus (IAV), which contains eight segments, and bunyaviruses (such as severe fever with thrombocytopenia syndrome virus (SFTSV)), which contains a tri-segment, *Mammarenavirus* utilizes the cap-snatching mechanism to start viral mRNA transcription (3–6). Cap-snatching involves recognizing capped cellular mRNAs followed by the cleavage of 10–14 nucleotides downstream by the polymerase’s endonuclease (EN) to provide a primer for viral mRNA transcription (7–10). The N-terminal 173-aa region of the LASV L protein has been identified as the EN domain, and its structure is highly homologous to other known viral endonucleases (10).

To date, no vaccines or specific antiviral agents against LASV have been developed. Therapeutic strategies are limited to ribavirin administration in the early stages of the illness. We have focused on developing entry inhibitors against LASV (11–13). Fragment-based drug discovery (FBDD) can identify early lead candidates for treatment, provide a backbone for drug optimization, and facilitate elucidation of the mechanism underlying the infection (14). As the EN domain might be an attractive target for replication inhibitors, shedding light on other negatively segmented RNA viruses, we used the LASV minigenome (MG) system to screen a library of 1,015 fragment-based drugs. After two rounds of screening, two benzotriazole compounds (F1204: 2-Amino-6-ethoxybenzothiazole, F1781: 2-Amino-6-chlorobenzothiazole) and 4,4’-dihydroxybiphenyl (F1597) were found to inhibit LASV MG activity. Furthermore, both benzotriazole compounds were capable of inhibiting other authentic negative segmented RNA viruses, such as LCMV and SFTSV.

## Results

### Screening of an FBDD library for inhibitors against LASV MG activities

To construct a high-throughput system to study viral replication in BSL-2 containment and facilitate high-throughput screening (HTS), LASV MG system based on the ambisense LASV S segment genome, in which the NP and GPC coding sequences were replaced with ZsGreen (ZsG) and *Gaussia princeps* luciferase (gLuc) reporter genes, co-transfected with the supportive pCAGGS-NP and pCAGGS-L plasmids, was constructed as previously reported (14). A ratio of 3:1:5 for *S* (ZsG/gLuc), pCAGGS-L, and pCAGGS-NP yielded an optimal signal-to-noise ratio >200 (Fig. 1A). Using ribavirin as the positive control, the Z′ factor was determined to be 0.87, indicating that the assay is robust and suitable for HTS.

**Fig. 1.**
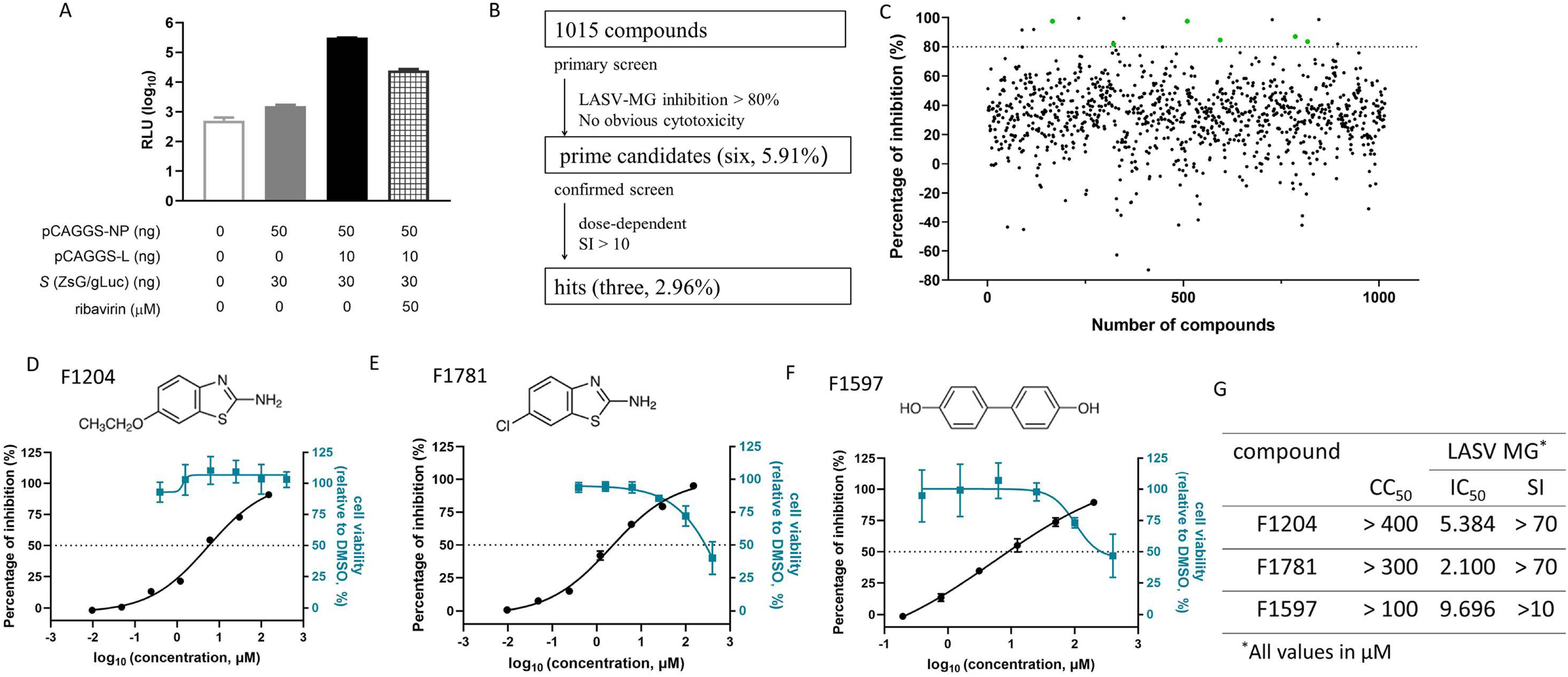
High-throughput screening (HTS) for inhibitors against LASV MG replication from a fragment-based drug library. (A) Validation of the LASV MG. Varying ratios of the LASV MG containing plasmids (total plasmid amount: 90 ng/well in 96-well plates) were transfected into Vero cells. Ribavirin of 50 μM, as a positive control, was added into the cells 5 h post-transfection. The cell lysates were subjected to test the gLuc activities 24 h later. (B) HTS assay flowchart. (C) HTS of a library of 1,015 fragment compounds for prime candidates inhibiting LASV MG activities. Each dot represents the percent inhibition achieved with each compound at a concentration of 100 μM. Six green dots represents the prime candidates with inhibition >80% and no obvious cytotoxicity. (D-F) Dose-response curves of F1204 (D), F1781 (E), and F1597 (F). Cells transfected with the LASV MG containing plasmids were treated in duplicate with each compound at the indicated concentrations; 24 h later, cells lysates were subjected to test the luciferase activities. Cell viability was evaluated using MTT assay. (G) IC_50_, CC_50_, and SI values for the three candidates. Data are presented as means ± standard deviations (SDs) for three to six independent experiments.

A flowchart of the HTS is depicted in Figure 1B. Inhibitors were defined as prime candidates, with LASV MG inhibition >80% and no apparent cytotoxicity at a concentration of 100 μM. Of the 1,015 tested compounds, six (5.91%) were identified as prime candidates (Fig. 1C). The confirmed screen was then carried out over a broader concentration range from 9.6 nM to 150 μM. Three compounds (2.96%) were identified as candidates based on their dose-dependent inhibition and a selective index (SI) >10. Among the three candidates, both F1204 and F1781 were benzotriazole compounds, with IC_50_ values of 5.384 and 2.100 μM, respectively. As both compounds showed little or mild cytotoxicity, the SI values of both compounds were >70. The third candidate, F1597 (4,4’-dihydroxybiphenyl), also showed dose-dependent inhibition against LASV MG with an IC_50_ of 9.696 μM (Fig. 1D to 1G).

### Both F1204 and F1781 inhibited LCMV replication

To investigate the inhibitory effects of the candidates on an authentic *Mammarenavirus*, BSL-2 compatible LCMV was utilized. First, the inhibition of LCMV MG activity was verified using LCMV MG on 293T cells. As shown in Figures 2A to 2C, although the inhibition on LCMV MG was slightly lower than that on LASV MG, all the candidates could effectively inhibit the LCMV MG activities in a dose-dependent manner. Notably, as F1597 showed cytotoxicity in 293T cells with a CC_50_ <100 μM, it was eliminated for further research. The antiviral effects of the remaining two candidates were validated using LCMV strain Cl13. As shown in Figures 2D and 2E, both F1204 and F1781 inhibited the authentic LCMV infection with IC_50_ values of 13.43% and 27.12%, respectively, suggesting that both compounds inhibited *Mammarenavirus* infection by blocking the replication step.

**Fig. 2.**
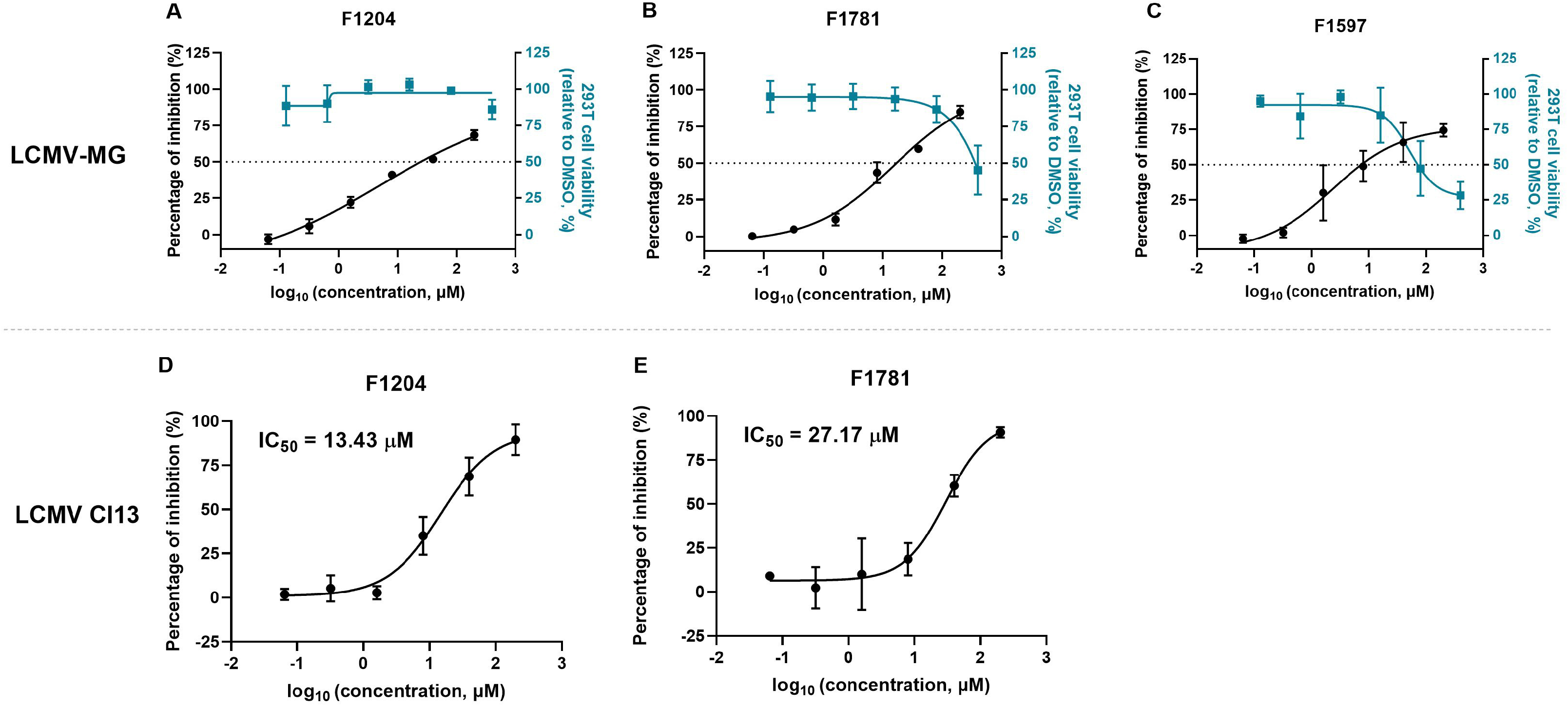
Inhibitory effects of the candidates against LCMV MG and the authentic LCMV Cl13 strain. (A-C) Inhibitory effects of F1204 (A), F1781 (B), and F1597 (C) against LCMV MG. LCMV MG containing plasmids (LCMV-*S*: 30 ng; LCMV-NP: 50 ng; LCMV-L: 10 ng) were transfected into 293T cells. Candidates with indicated concentration were added into the cells 5 h post-transfection. The cell lysates were subjected to test the gLuc activities 24 h later. The 293T cells viabilities were tested using MTT assay. (D-E) Dose-response curves of F1204 (D) and F1781 (E) against authentic LCMV Cl13 strain. Vero cells were treated with either compound; 1 h later, LCMV Cl13 of MOI: 0.1 was added to the cells; the supernatant was removed 1 h postinfection, and the cells were incubated with either compound for additional 23 h. The Vero cell lysates were assessed by qRT-PCR. Data are presented as means ± standard deviations (SDs) for three independent experiments.

### F1204 and F1781 had little effects on *Mammarenavirus* entry and budding

To further assess the role of both F1204 and F1781 on other steps in the life cycle of *Mammarenavirus*, the pseudotype virus was used to evaluate the effect on the virus entry step. In contrast, the Z protein-expressing plasmid was used to investigate viral budding. Figures 3A and 3B showed neither F1204 nor F1781 blocked LASVpv infection as the percentage of inhibition barely reached 50%, even at the highest tested concentration (200 μM). Similarly, both compounds could hardly block LCMVpv infection (Fig. 3C and 3D), suggesting that neither compound had any effect on *Mammarenavirus* entry. Furthermore, we investigated the effects of the hit compounds on the budding ability of the matrix Z protein by detecting virus-like particles (VLPs) with a Z self-budding assay (15–17). As shown in Figures 3E–3H, even when treated with either F1204 or F1781 at 100 μM, LASV Z could efficiently produce VLP, relative to the vehicle group, suggesting that neither F1204 nor F1781 impaired *Mammarenavirus* Z budding ability. Collectively, these results demonstrated that both F1204 and F1781 exerted antiviral effects on the replication of mammarenaviruses.

**Fig. 3.**
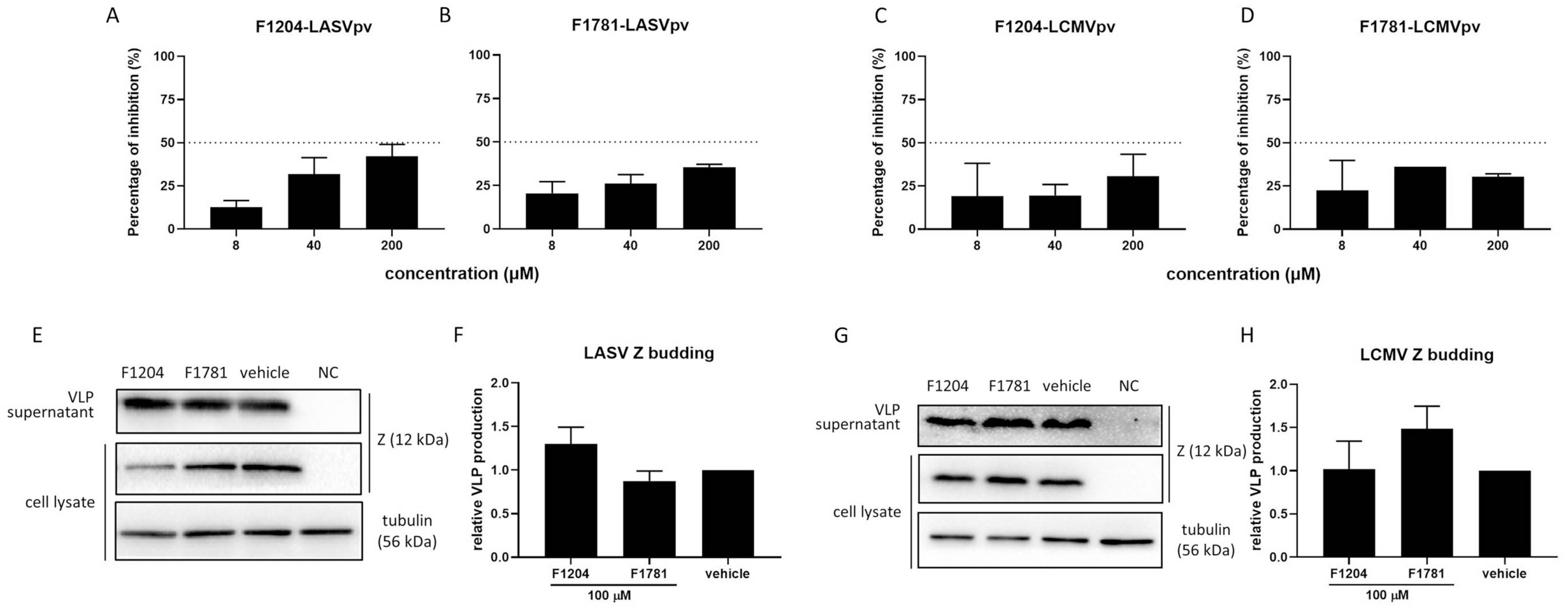
Effects of F1204 and F1781 on mammarenavirus entry and budding. (A and B) Effects of F1204 and F1781 on LASV entry. Vero cells were preincubated with F1204 (A) and F1781 (B), respectively, for 1 h, followed by incubation with LASVrv (MOI, 0.1) in the presence of compounds for 1 h. The cell lysates were assessed for luciferase activities 24 h post-infection. (C and D) Effects of F1204 and F1781 on LCMV entry. (E) Effects of F1204 and F1781 on LASV VLP production. 293T cells were transfected with pCAGGS-LASV Z; 4 h later, compounds of 100 μM were added and incubated for 48 h. The supernatant and cell lysate were collected for WB assay. (F) Quantification results of WB assay were presented as the mean ± SD from five independent experiments.

### Binding ability of both hits to LASV EN

To investigate whether the endonuclease was the target of both hit compounds, we further analyzed the binding affinity of the hits to LASV EN using isothermal titration calorimetry (ITC) in the presence of divalent metal ions (Mg^2+^ and Mn^2+^). As shown in Figure 4, F1204 and F1781 bound to LASV EN with the equilibrium dissociation constant (*K*_*D*_) of 35.3 and 12.8 μM, respectively, suggesting that LASV EN might serve as the target of both F1204 and F1781.

**Fig. 4.**
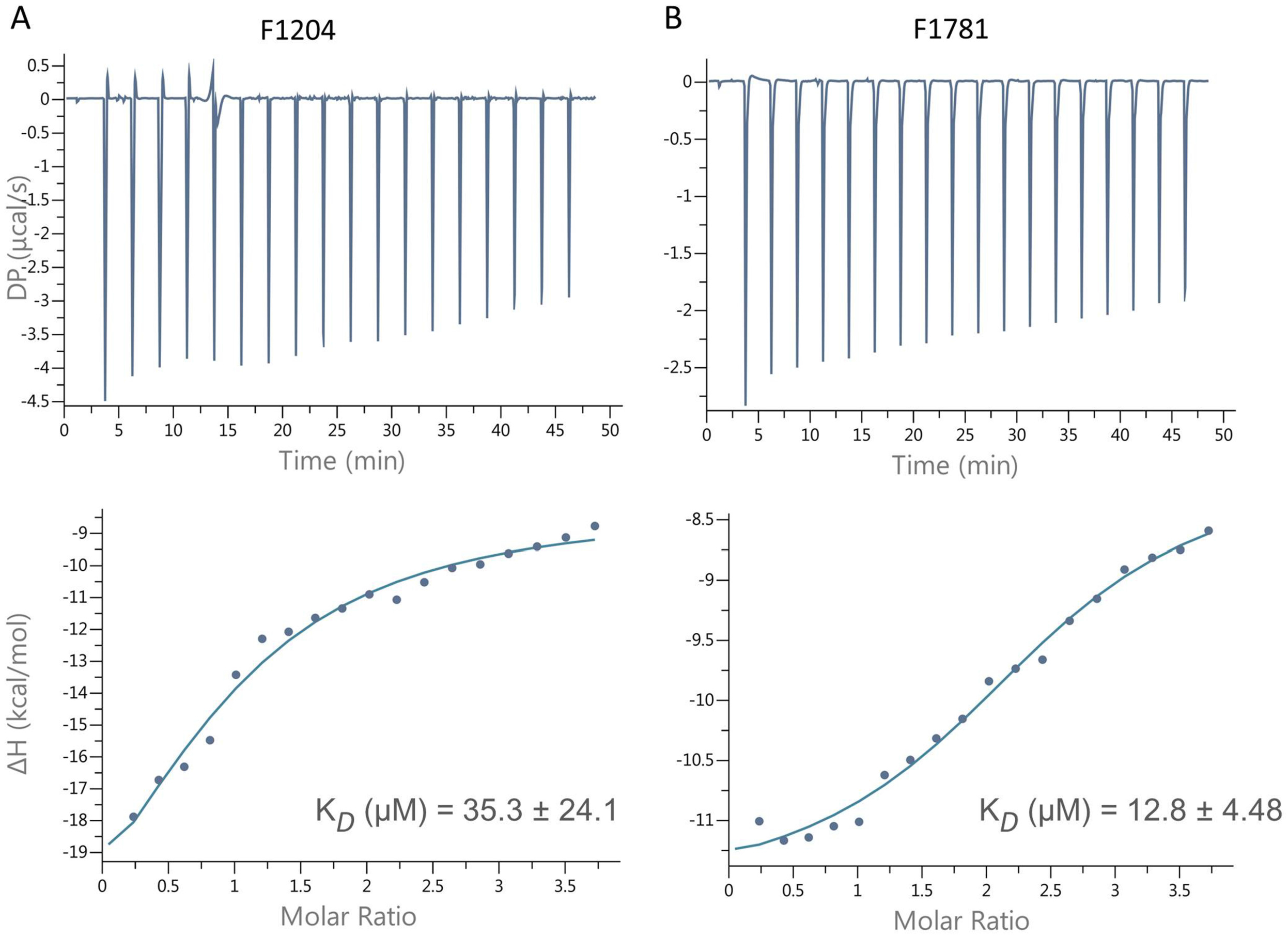
Binding of F1204 and F1781 to LASV endonuclease measured by ITC. F1204 (A) and F1781 (B) binding to 45 μM LASV endonuclease protein were measured at 25 °C by ITC. The upper plot showed the binding isotherm and the lower plot shows the integrated values.

### Combinatorial effects of hit compounds and *Mammarenavirus* entry inhibitor

As both F1204 and F1781 targeted the endonuclease and thus inhibited *Mammarenavirus* replication, we hypothesized that these replication inhibitors might synergize with the entry inhibitor. To address this, we utilized casticin, a botanical drug demonstrated to inhibit *Mammarenavirus* entry by blocking GPC-mediated membrane fusion (13), to study the combined therapeutic effects. The prototype *Mammarenavirus*, LCMV, was used again because each step of the authentic virus life cycle could be studied in the BSL-2 lab. As expected, both the combinations of F1204 with casticin and F1781 with casticin had significant synergistic antiviral effects (18, 19) with different volumes of 194.54 and 148.45 nM^2^ %, respectively (Fig. 5) (20).

**Fig. 5.**
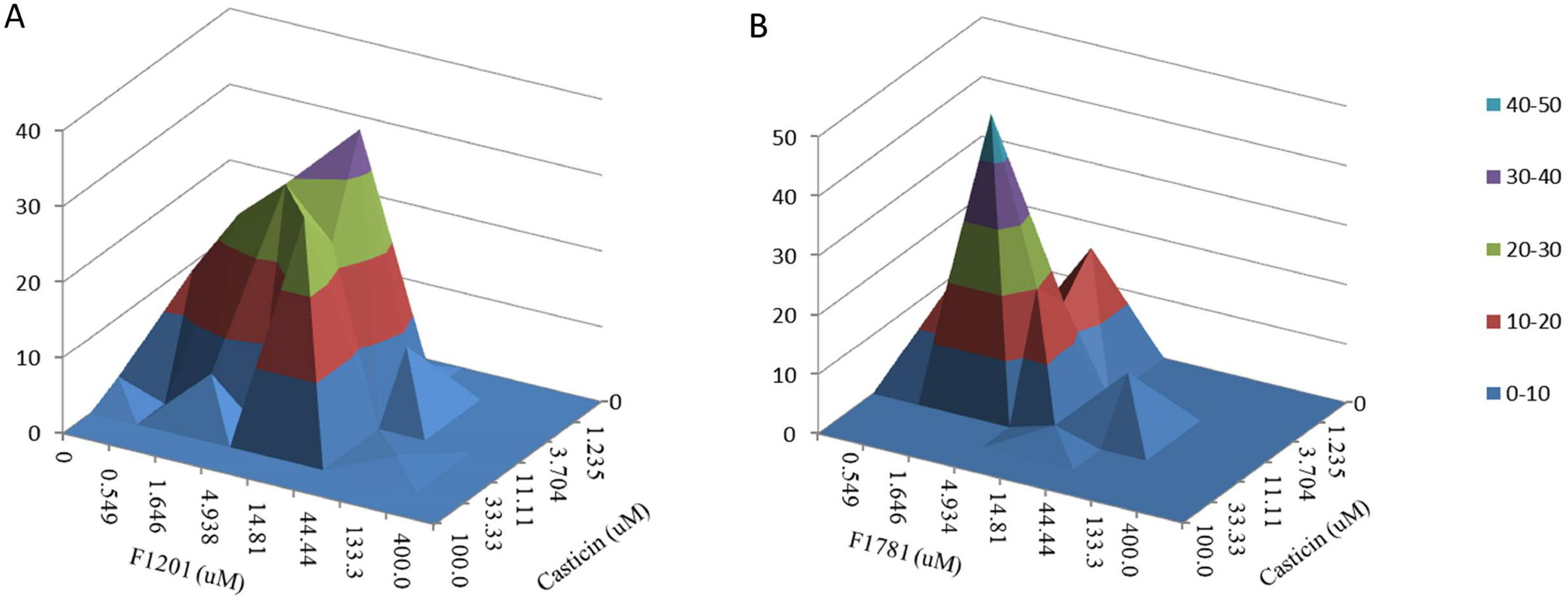
Drug-drug interactions of F1204 and F1781 with casticin. (A and B) Differential surface plots at 95% confidence level (CI) were calculated and generated using MacSynergy II for the drug-drug interactions for evaluating combinations of F1204 (A) and F1781 (B) with casticin targeting LCMV.

### F1204 and F1781 exerted an antiviral effect against SFTSV

To test whether both hit compounds could extend the antiviral spectrum to other negative segmented RNA viruses, we investigated the inhibitory effect of both compounds against SFTSV, a tick-borne virus belonging to the genus *Banyangvirus* (family Phenuiviridae) (21). As shown in Figure 6, both F1204 and F1781 inhibited SFTSV infection in a dose-dependent manner. However, the inhibitory effect was less pronounced than that of LCMV. F1204 and F1781 (200 μM) robustly inhibited LCMV infection to ~90% (Fig. 2), while the inhibition of SFTSV was only ~60% (Fig. 6), which was similar to a previous report that the approved IAV EN inhibitor, L-742001, which inhibited SFTSV only at high concentrations (>0 μM) (5). This might be due to the subtle structural difference in the EN domain, which resulted in the difference in the antiviral effects (6).

**Fig. 6.**
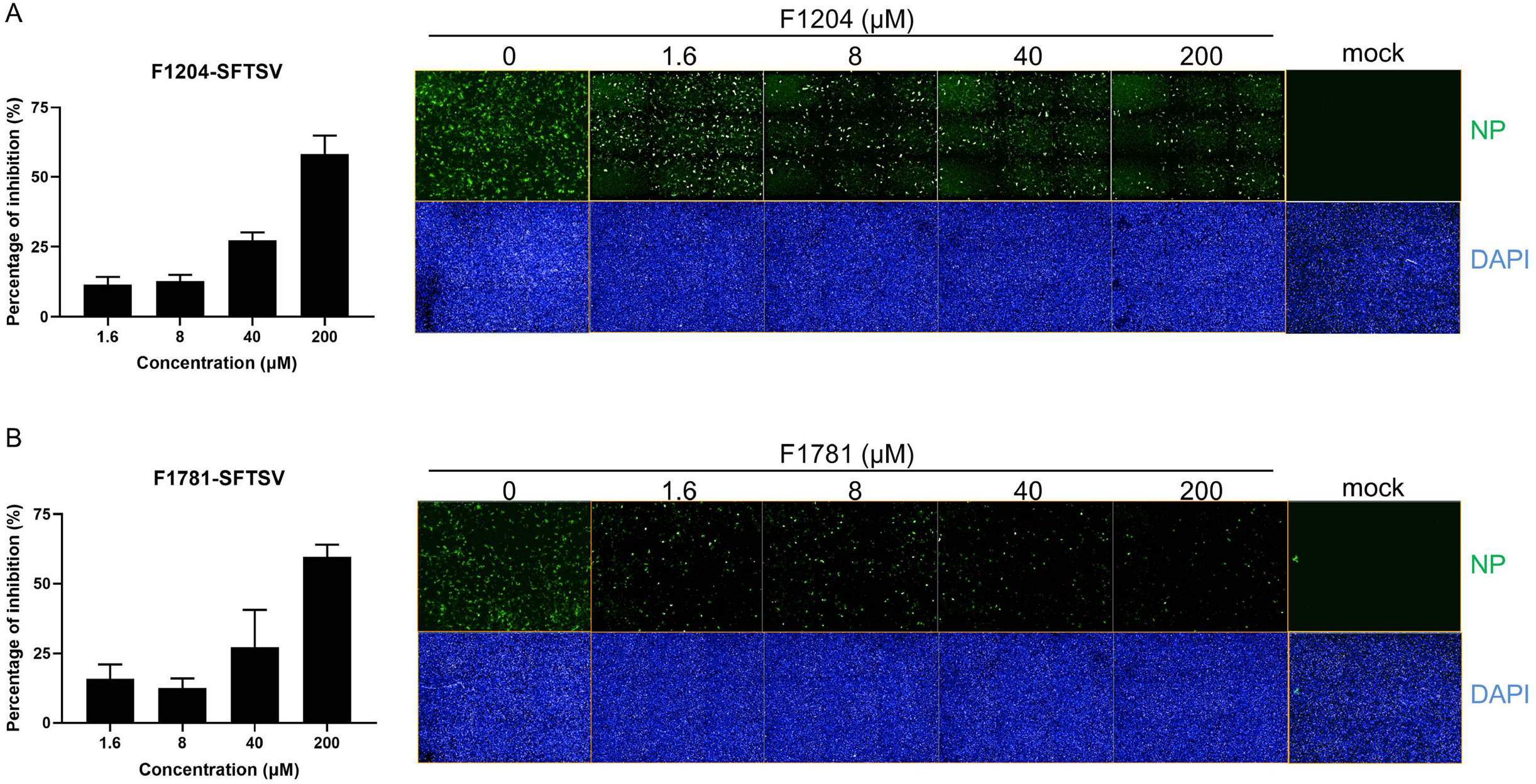
Inhibitory effects of F1204 and F1781 against SFTSV. (A) Inhibitory effect of F1204 against SFTSV. Vero cells were treated with F1204; 1 h later, SFTSV of MOI: 0.1 was added to the cells; the supernatant was removed 1 h postinfection, and the cells were incubated with F1204 for additional 23 h. The percentage of inhibition was calculated using the Harmony 3.5 software in an Operetta high-content imaging system (PerkinElmer) (left), and the cells were imaged using the same system (right). (B) Inhibitory effect of F1781 against SFTSV. Data are presented as means ± standard deviations (SDs) for three to four independent experiments.

## Discussion

Based on the 2020 International Committee on Taxonomy of Viruses (ICTV)-taxonomic update reports, the genus *Mammarenavirus*, family *Arenaviridae*, is added to the order Bunyavirales, which belongs to the class Ellioviricetes, subphylum Polyploviricotina. Polyploviricotina contains a negative-sense RNA virus that encodes the L protein without mRNA capping activity. The other class in Polyploviricotina is Insthoviricetes, which includes the family Orthomyxoviridae, which contains IAV and IBV (1). The cap-snatching mechanism in the initial phase of mRNA transcription employed in all negative-sense, segmented RNA viruses is a potential target for developing broad-spectrum antiviral drugs. Notably, the endonuclease inhibitor baloxavir marboxil (BXM) was approved in 2018 to treat IAV and IBV infections (22). The active metabolite (baloxavir acid, BXA) of BXM targets the IAV EN with a metal chelating mechanism (23–25), similar to the first-generation endonuclease inhibitors, such as L-742,001 (26, 27). It has been recently reported that BXA exhibited a micromolar level inhibition on SFTSV EN activity and SFTSV plaque formation (5).

As cap-snatching is the replication strategy shared by all segmented negative-sense RNA viruses, and these viruses have a similar EN structure, EN is an attractive target for small molecule inhibitors. In this study, we utilized the FBDD library to screen for inhibitors targeting LASV EN. Among the three identified candidates, F1204 (2-Amino-6-ethoxybenzothiazole) and F1781 (2-Amino-6-chlorobenzothiazole) are benzotriazole compounds. To this end, we first reviewed all the nine benzotriazole compounds included in the current library. We found that three additional compounds, F1145, F1406, and F1486, showed inhibition levels of 50%–79% on the primary screen, while the other four compounds showed cytotoxicity or little inhibition (Table 1). Based on these results, we hypothesized that benzotriazole might serve as a backbone for the development of hit drugs. We selected two approved drugs, riluzole: 2-Amino-6-(trifluoromethoxy) benzothiazole, and Frentizole: 1-(6-Methoxy-2-benzothiazolyl)-3-phenylurea with the benzotriazole backbone and tested its effects on LASV MG. Unfortunately, neither showed inhibition of LASV MG activity. These results, however, were instrumental in highlighting the structure-activity relationship optimization of lead compounds.

**Table 1.**
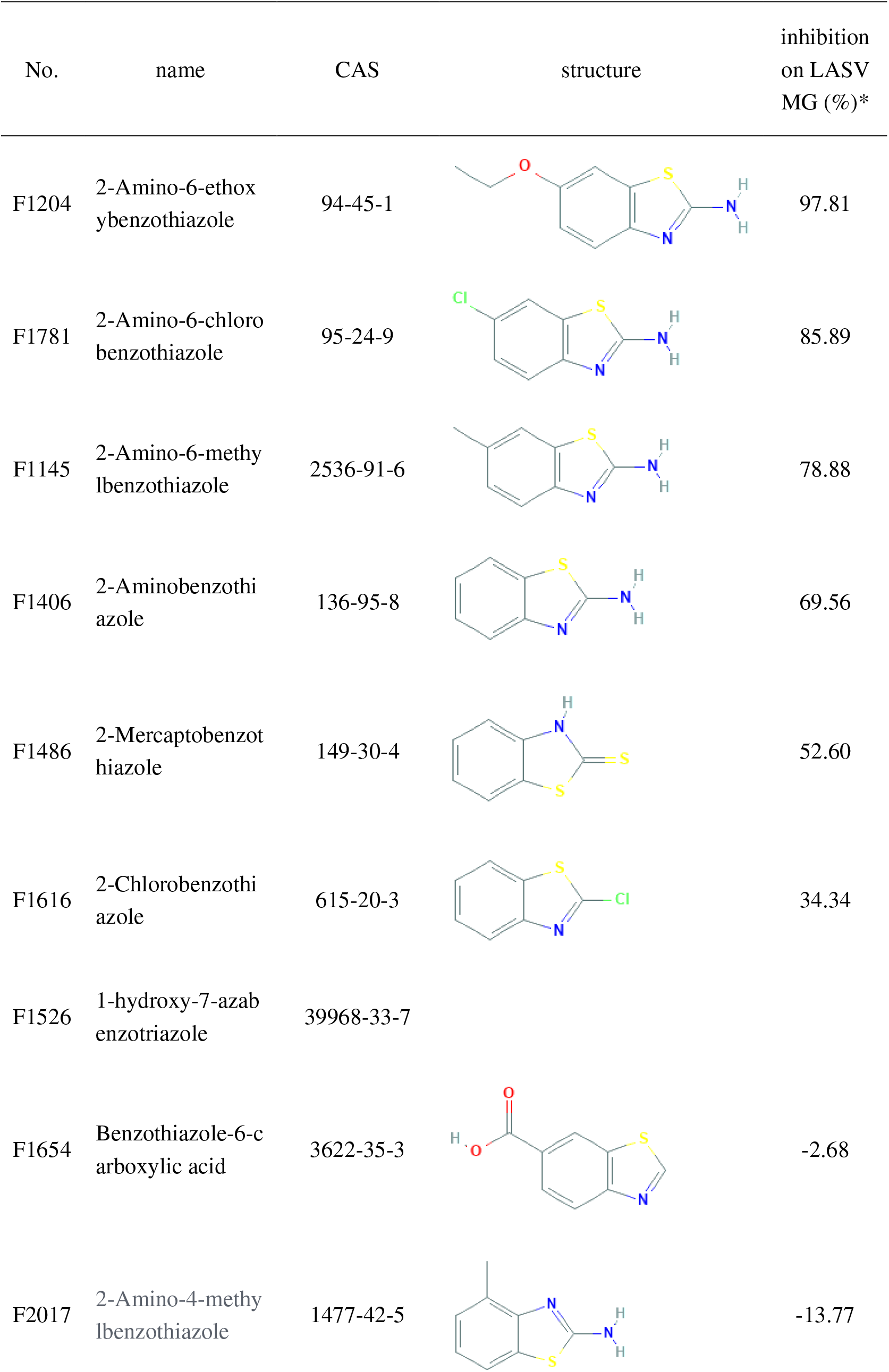
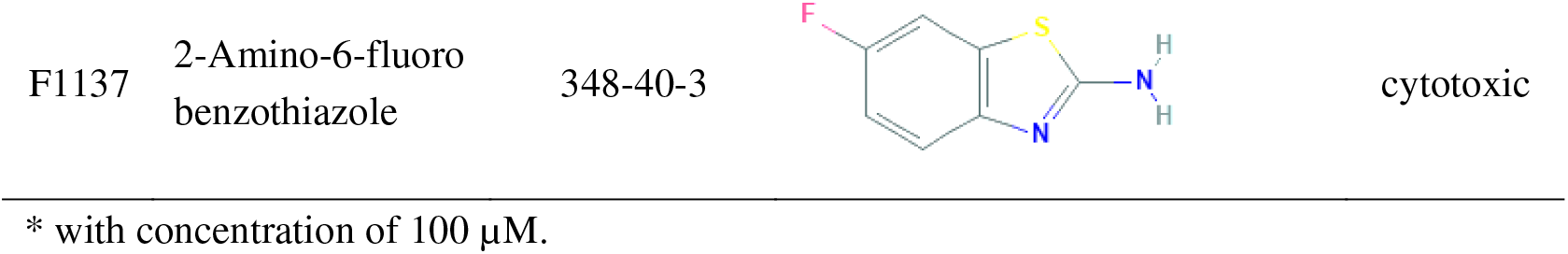
Benzotriazole compounds in FBDD library.

The FBDD library contains compounds that are small in size and low in molecular weight (~200 Da), and FBDD library screens usually utilize biophysical methods such as X-ray crystallography, nuclear magnetic resonance, and mass spectrometry to identify the hits (28, 29). The binding affinity of the hits to the targeting protein is usually weak because the fragments could only occupy part of the binding pocket. Sometimes, the hit fragments showed no activity in the functional assays (30). We screened the FBDD library using the enzyme activity-based high concentration screening strategy, which has been reported to be an effective and rapid approach for identifying fragments (31, 32). Both the hit compounds, F1204 and F1781, showed the micromolar binding ability to LASV EN affinity. Furthermore, both hit compounds could effectively inhibit authentic LCMV and SFTSV infection, suggesting the potential to develop hit compounds to broad-spectrum antiviral inhibitors. The synergistic effects of the combination of the hit compounds with the entry inhibitors shed light on developing a successful treatment for pathogenic segmented, negative-sense RNA virus infectious diseases.

## Materials and Methods

### Cells and viruses

Vero, 293T, BSR-T7, and BHK-21 cells were cultured in Dulbecco’s modified Eagle’s medium (DMEM; HyClone, Logan, UT, USA) supplemented with 10% fetal bovine serum (Gibco, Grand Island, NY, USA).

LCMV clone 13 was rescued using genome RNA L (DQ361066) and S (DQ361065) segments, as previously reported (13, 33). Briefly, BSR-T7 cells were transfected with a mixture of plasmids containing pCAGGS-LCMVNP (300 ng), pCAGGS-LCMV-L (600 ng), pT7-LCMV-L (600 ng), and pT7-LCMV-S (300 ng). The supernatant was collected 72 h later and inoculated into BHK-21 cells for amplification. The titer of LCMV was determined to be

1 × 10^7^/ml by plaque assay. SFTSV (WCH/97/HN_Henan_2011) was provided by the Microorganisms and Viruses Culture Collection Center, Wuhan Institute of Virology, Chinese Academy of Sciences. The titer of SFTSV was determined to be 1 × 10^5^/ml by the TCID_50_ assay. LASVpv and LCMVpv were generated using the VSV-based pseudotype virus system (34, 35). Briefly, 293T cells transfected with pCAGGSGPC were infected with pseudotype VSV, in which the G gene was replaced with a luciferase gene. The culture supernatants were harvested 24 h later, and the viral titers of LASVpv and LCMVpv were determined to be 3 × 10^7^/ml and 1 × 10^6^/ml, respectively (11, 13).

### MG assay

For the LASV MG assay, Vero cells were seeded at 1.5 × 10^4^ cells per well in a 96-well plate. After overnight incubation, cells were transfected with 90 ng of mixed plasmids with a 3:1:5 ratio of pLASV-S (ZsG/gLuc), pCAGGS-L, and pCAGGS-NP. The minigenome is based on the LASV S segment, as previously reported (14). The NP and GPC were replaced with ZsG and gLuc, respectively, and the authentic untranslated regions intergenic regions were unchanged (Josiah strain; GenBank number HQ688673.1). The cells were lysed 24 h later, and luciferase activity was measured using the Rluc assay system (Promega, Madison, WI). For the LCMV MG assay, 293T cells were used.

### HTS assay of an FBDD library

A library of 1,015 fragment-based drugs was purchased from Selleck Chemicals (Cat: L1600; Houston, TX, USA). Compounds were stored in 10 mM stock solution in DMSO at −80 °C until use. The first-round HTS was performed, as shown in Figure 1B. Briefly, cells transfected with MG mixed plasmids were treated in duplicate with the compounds (100 μM); 24 h later, cells were lysed to measure luciferase activity. Prime candidates were identified using criteria of no apparent cytotoxicity and an average >80% inhibition in duplicate wells and subsequently confirmed by serial dilution in triplicate plates to evaluate the IC_50_ (GraphPad Prism 6). Using the criteria of dose-dependent inhibition and SI > 10, three compounds were selected. Cytotoxicity was determined using the 3-(4,5-dimethyl-2-thiazolyl)-2,5-diphenyl-2H-tetrazolium bromide (MTT) assay.

### Antiviral assay

Vero cells were seeded at a density of 1 × 10^4^ cells per well in a 96-well plate. After incubation overnight, cells were incubated with compounds at the indicated concentrations for 1 h. LCMV and SFTSV (MOI: 0.1) were added to the cells and incubated for 1 h. The cells were then incubated with the compounds for an additional 24 h. For the anti-LCMV assay, the cell lysates were subjected to RT-qPCR using the primers LCMV-f: AGAATCCAGGTGGTTATTGCC and LCMV-r: GTTGTAGTCAATTAGTCGCAGC and GAPDH-f: 5’-TCCTTGGAGGCCATGTGGGCCAT-3’ and GAPDH-r: 5’-TGATGACATCAAGAAGGTGGTGAAG-3’. In the anti-SFTSV assay, the cells were subjected to an immunofluorescence assay using rabbit anti-NP serum (kindly provided by Prof. Fei Deng at Wuhan Institute of Virology, CAS).

### VLP assay

The VLP assay was conducted as described previously (15–17). Briefly, 293T cells were seeded at 8 × 10^5^ cells per well in a 6-well plate. After overnight incubation, the cells were transfected with pCAGGS-LASV-Z and pCAGGS-LCMV-Z. Four hours later, 100 μM hit compounds were added to the cells for 48 h. The supernatant was centrifuged at 15,000 *× g* for 10 min at 4 °C. The supernatant was then treated with 8% PEG8000 and 0.5 M sodium chloride, followed by centrifugation at 15,000 *× g* for 15 min. The samples collected from the supernatant and cell lysate were subjected to WB assay using anti-LASV Z antibody (GeneTex, cat: GTX134874, CA, USA) and anti-HA antibody (Proteintech, cat: 66006-2-Ig, IL, USA) to detect the LASV and LCMV VLPs, respectively.

### LASV EN protein expression and purification

*E. coli* codon-optimized coding sequences were synthesized (Sanger) for residues 1-173 (LASV EN) of LASV L protein (7). LASV EN was cloned into pET-9a (Novagen) with an N-terminal His-tag, and a tobacco etch virus (TEV) cleavage site (MGHHHHHHDYDIPTTENLYFQG-). The protein was expressed in *E. coli* strain BL21 (DE3) at 16 °C in LB media for 16 h after induction with (IPTG 0.2 mM). The protein was purified using a Ni-NTA column followed by removing the His-tag by TEV protease as previously described (6).

### ITC assay

ITC assays were performed at 25 °C using a MicroCal PEAQ-ITC (MicroCal, Inc.). The experiments comprised 19 injections of 2 μl of 0.9 mM compound into the sample cell containing 200 μl of 45 μM Lassa EN. The heat produced by the compound dilution in the buffer was subtracted from the heat obtained in the presence of the protein. Binding isotherms were fitted to one-site binding using MicroCal PEAQ-ITC Analysis software.

## ACKNOWLEDGMENTS

We thank Dr. César G. Albariño for general advice on the creation of the LASV and LCMV minigenome system. We thank Dr. Liangdan Fei from Wuhan University for providing assistance in the ITC assay. We thank the Center for Instrumental Analysis and Metrology and Core Facility and Technical Support, Wuhan Institute of Virology, for providing technical assistance.

This work was supported by the National Key Research and Development Program of China (2018YFA0507204), the National Natural Sciences Foundation of China (31670165), Wuhan National Biosafety Laboratory, Chinese Academy of Sciences Advanced Customer Cultivation Project (2019ACCP-MS03), and the Open Research Fund Program of the State Key Laboratory of Virology of China (2018IOV001).

